# Low-dose VSV-EBOV vaccination provides rapid protection from lethal Ebola virus challenge

**DOI:** 10.64898/2026.02.14.705917

**Authors:** Andrea Marzi, Wakako Furuyama, Amanda J. Griffin, Friederike Feldmann, Kyle W. Shifflett, Elizabeth R. Wrobel, Kyle L. O’Donnell, Patrick W. Hanley, Heinz Feldmann

**Affiliations:** Laboratory of Virology, Division of Intramural Research, National Institute of Allergy and Infectious Diseases, National Institutes of Health, Hamilton, MT 59840, USA; Rocky Mountain Veterinary Branch, Division of Intramural Research, National Institute of Allergy and Infectious Diseases, National Institutes of Health, Hamilton, MT 59840, USA

**Keywords:** Filovirus, vesicular stomatitis virus, immunity, rapid protection

## Abstract

In the decade since the West African Ebola virus (EBOV) epidemic, several medical countermeasures against this often-fatal hemorrhagic disease have been approved by regulatory authorities for human use. This includes monoclonal antibody-based therapies and vaccines which have been stockpiled in limited quantities. One of the vaccines is based on vesicular stomatitis virus (VSV) expressing the EBOV glycoprotein (GP) as the immunogen. The vaccine is stockpiled in limited quantity for emergency use. A single high dose has been shown to rapidly protect humans within 10 days. We developed an updated version of this vaccine expressing the GP from the 2015 EBOV-Makona isolate. Here, we wanted to determine the protective efficacy within 10 days of a single moderate dose (10-fold and 1,000-fold dilution) of the updated vaccine in nonhuman primates (NHPs). As a comparator we included a 1000-fold dilution dose group of the approved vaccine expressing the EBOV-Kikwit GP. While we achieved uniform protection with the approved vaccine at the moderate dose, only 50% of the NHPs receiving the same dose of the updated vaccine expressing the EBOV-Makona GP were protected. This study highlights the importance of evaluating VSV-based vaccine stocks expressing different filovirus GPs in preclinical models prior to progression with clinical development. Our study also highlights that rapid vaccination with reduced doses still leads to protection but at the cost of “sterile” immunity raising concerns regarding EBOV persistence and potential downstream transmission. Therefore, lower vaccine doses should only be considered in cases of severe vaccine shortage.

## Introduction

Ebola virus (EBOV), member of the *Filoviridae* family, is a highly pathogenic virus and of eminent concern to public health in endemic areas in Africa ^1^. Since the 2013-2016 EBOV epidemic in West Africa, annual outbreaks have occurred mainly in the Democratic Republic of the Congo (DRC)^2^. EBOV infection causes Ebola virus disease (EVD) that results in fatalities in up to 90% of patients if untreated ^3^. In 2019, the European Medicines Agency and the United States Food and Drug Administration (FDA) approved the use of EBOV vaccines, which several African countries have subsequently used in recent outbreaks ^4^. In addition, there are also two monoclonal antibody-based treatments approved for the use against EVD ^5^.

One of the vaccines is a live-attenuated vesicular stomatitis virus (VSV)-based vector expressing the EBOV-Kikwit glycoprotein (GP) instead of its native glycoprotein (G) (VSV-EBOV_Kik_) ^6^. This VSV-EBOV vaccine (also known as rVSV-ZEBOV and Ervebo) protects nonhuman primates (NHPs) after a single high dose vaccination within 7 days ^7,8^ and has been successfully used in the DRC during the latest outbreaks ^9,10^. A human clinical trial conducted in Guinea during the 2013-2016 EBOV epidemic and data from more recent outbreaks in the DRC confirmed the fast-acting property of this vaccine in humans; it was protective within 10 days after administration ^10,11^. In response to the large West African EBOV epidemic we have updated VSV-EBOV to express the EBOV-Makona GP (VSV-EBOV_Mak_) and demonstrated that a dose as low as 10 plaque-forming units (PFU) uniformly protects NHPs from lethal disease 4 weeks after vaccination ^12^. Due to previous vaccine shortages for outbreak response and limitations in production and stockpiling ^13^, we wanted to investigate if a lower dose of VSV-EBOV_Mak_ could still protect rapidly from lethal disease overcoming potential shortcomings in vaccine availability during future outbreaks.

We evaluated 2 vaccine doses, 1×10^6^ and 1×10^4^ PFU (10-fold or 1,000-fold dilution from original dose, respectively), of VSV-EBOV_Mak_ in comparison to 1×10^4^ PFU of VSV-EBOV_Kik_, the licensed vaccine. A single vaccine dose was administered to NHPs 10 days before the lethal challenge mimicking the successful human clinical trials ^10,11^. We observed uniform protection with 1×10^6^ PFU VSV-EBOV_Mak_ and 1×10^4^ PFU VSV-EBOV_Kik_; in contrast, 50% of the NHPs succumbed to the disease when vaccinated with 1×10^4^ PFU VSV-EBOV_Mak_.

## Results

### Single low-dose vaccination with VSV-EBOV is protective within 10 days

In a previous study we demonstrated protection after a single vaccination with a low-dose of VSV-EBOV_Mak_ against EBOV-Makona challenge in NHPs ^12^. Here, we expanded upon this data and evaluated the fast-acting potential of reduced doses of VSV-EBOV_Mak_ vaccination against EBOV-Makona challenge in NHPs. We included vaccination with a single lower dose of VSV-EBOV_Kik_ ^8,14^ to investigate the efficacy of an “unmatched” vaccine to the challenge virus. We compared 1×10^6^ and 1×10^4^ PFU of the VSV-EBOV_Mak_ to 1×10^4^ PFU VSV-EBOV_Kik_ in groups of 4 NHPs each.

Positive and negative control groups were included as well: two NHPs received 1x 10^7^ PFU VSV-EBOV_Mak_, a dose previously shown to protect NHPs from lethal disease within 7 days ^8^, and 4 NHPs received 1x 10^7^ PFU VSV-MARV, a control vaccine (Fig. S1A). Vaccination occurred on day - 10 by the intramuscular (IM) route and was followed by physical exams including blood draws (Fig. S1B). Unfortunately, one control NHPs succumbed prior to EBOV challenge to complications from anesthesia due to a previously unidentified cardiac condition. This animal was removed from the study. After vaccination, VSV RNA levels in the blood were monitored and showed that vaccine RNA levels were maintained similarly between the groups. However, a significantly higher level of VSV RNA was measured 3 days post-vaccination (day -7) for the 10^7^ PFU VSV-EBOV_Mak_, 10^4^ PFU VSV-EBOV_Kik_ and control groups (10^7^ PFU VSV-MARV) (Fig. S1C). All NHPs were infected IM on day 0 with 1,000 PFU of EBOV-Makona. The animals were closely monitored throughout the study, and a daily clinical score was calculated based on approved parameters. Only the control group and 2 NHPs vaccinated with 10^4^ PFU VSV-EBOV_Mak_ developed severe clinical signs of EVD (Fig. 1A) and were euthanized 5, 6 or 8 days post-challenge (DPC) (Fig. 1B). NHPs vaccinated with 10^4^ PFU VSV-EBOV_Kik_ or higher doses of VSV-EBOV_Mak_ displayed only transient mild signs of the disease (ruffled appearance and decreased appetite) and recovered quickly (Fig. 1A). EBOV was only isolated from the blood of NHPs that succumbed to infection during the acute phase of the disease (Fig. 1C). Our previous study had shown a correlation between EBOV sGP levels and viremia levels in infected NHPs ^15^. This study confirmed this observation as EBOV sGP was only present in the serum of viremic NHPs euthanized during the acute phase of disease (Fig. 1D). Complete blood counts revealed that only those NHPs who succumbed to challenge developed thrombocytopenia (Fig. 1E), a characteristic sign of EVD ^1,3^. In addition, EBOV was only isolated from tissue samples collected at necropsy form NHPs that succumbed to disease (Fig. 1F). Similarly, serum chemistry values showed elevated levels of liver transaminases (ALT, AST; Fig. S2A,B) and alkaline phosphatase (ALP; Fig. S2C) which have been reported in NHPs that succumbed during acute disease indicating liver pathology^8,12^. Blood urea nitrogen (BUN) was elevated in all NHPs that exhibited acute EVD (Fig. S2D) and elevations in creatinine were observed in some EVD-affected NHPs (Fig. S2E). These values are indicative of either severe dehydration or renal dysfunction; however, additional testing is necessary to distinguish which was not conducted. These values match other reports of NHPs that succumb to EVD^1,3^. All vaccinated and protected NHPs had little if any changes in blood cell counts and serum chemistry values (Fig. 1F, S2A-E).

**Figure 1.**
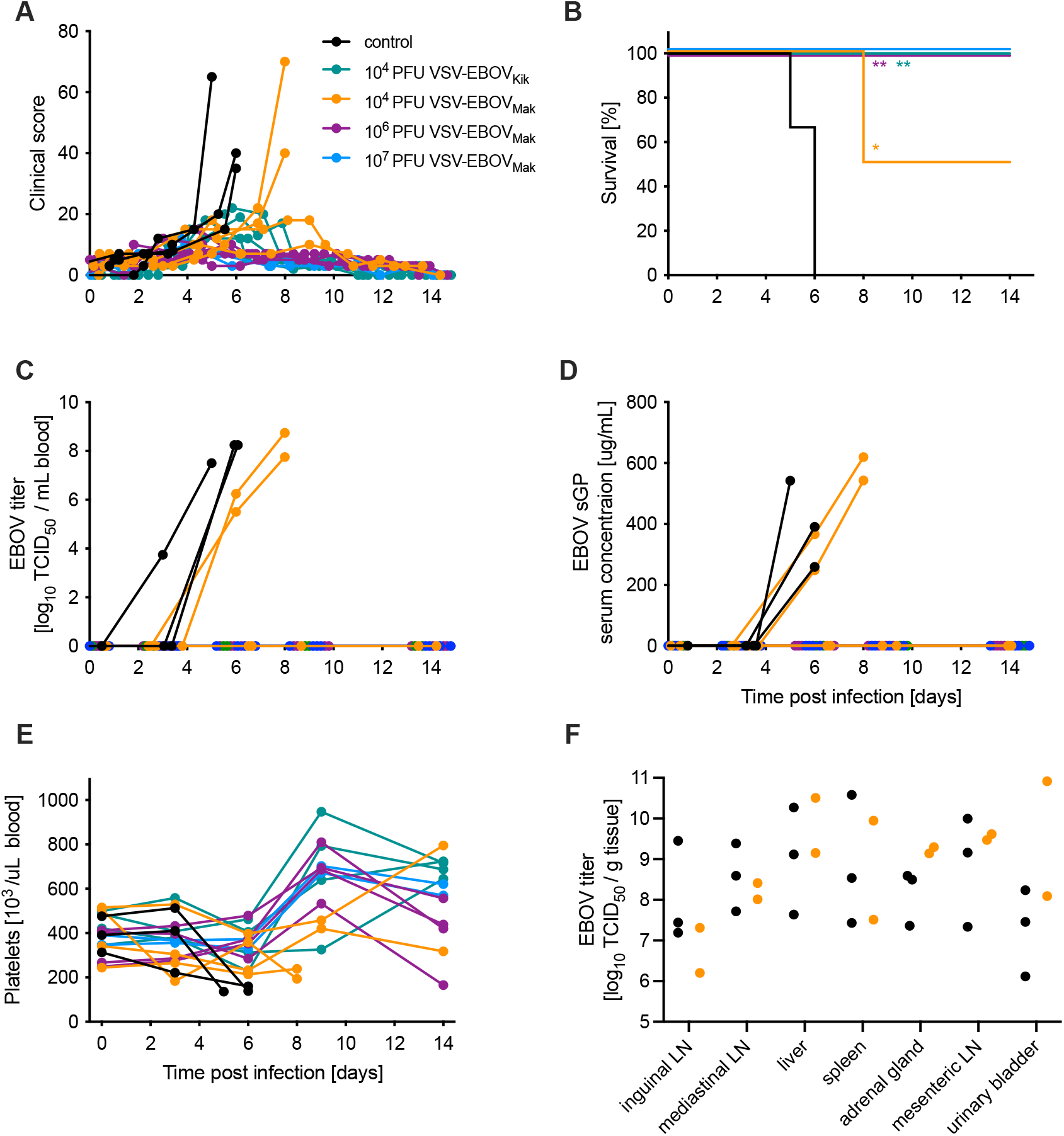
Survival and clinical parameters of EBOV-infected NHPs. Cynomolgus macaques were IM-vaccinated with respective VSV vaccines and challenged with EBOV 10 days later. **(A)** Clinical scores, **(B)** survival curves, **(C)** EBOV blood titers are shown for the critical phase of the disease. **(D)** EBOV sGP levels in serum samples over time. **(E)** Platelet counts in whole blood over time. **(F)** EBOV titers in key target tissues are depicted for all animals that reached the humane endpoint due to the infection. Statistical significance of vaccine groups versus controls is indicated (* *p*< 0.05, ** *p*<0.01).

### Only non-protected NHPs develop a “cytokine storm”

Because lethal EVD has been associated with a dysregulated cytokine response known as “cytokine storm” ^1,3^, we analyzed serum cytokine levels in the control and low-dose vaccine groups. As we only had two NHPs in the positive control group (10^7^ PFU VSV-EBOV_Mak_) we could not expect to obtain statistically significant differences and omitted serum cytokine analysis for this group. In concordance with previous data ^8,16^ we observed high levels of cytokines and chemokines at the time of euthanasia 5-8 DPC in the five NHPs that succumbed to disease (Fig. 2). In response to the VSV vaccination, IFNα was upregulated on -9 DPC in all NHPs and returned to base line by -7 DPC (Fig. 2). At 24 hours post vaccination (-9 DPC) IL-1ra, IL-10, and MCP-1 were increased with highest levels in the 10^4^ PFU VSV-EBOV_Kik_ group (Fig. 2). Interestingly, at the time of challenge (0 DPC) NHPs in the 10^4^ PFU VSV-EBOV_Mak_ group demonstrated significantly lower levels of IL-1ra and MCP-1 compared to the 10^4^ PFU VSV-EBOV_Kik_ group. Other cytokines and chemokines did not result in significant differences at any time point samples were obtained (Fig. 2).

**Figure 2.**
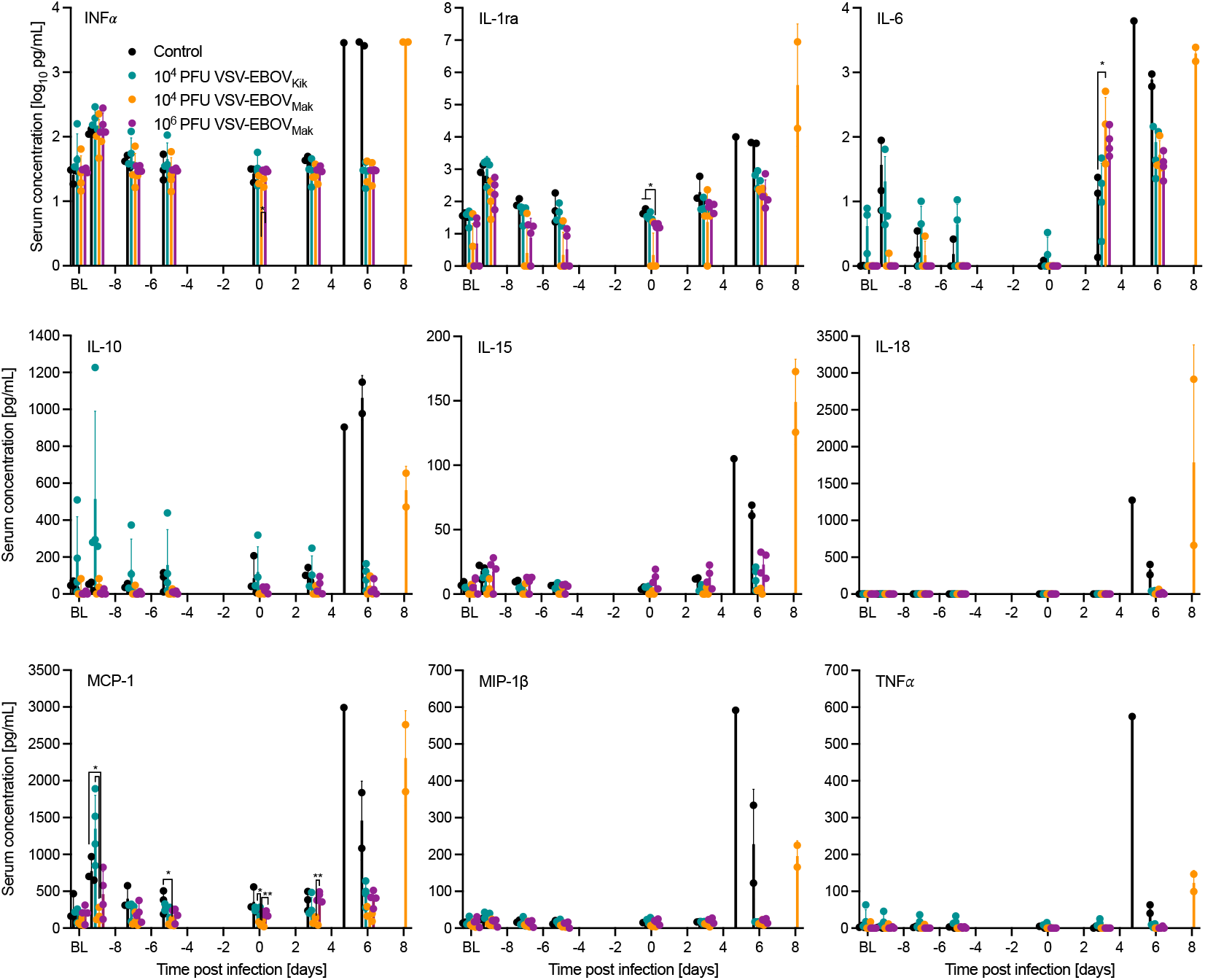
Cytokine responses in vaccinated and challenged NHPs. Serum concentrations for INFα, IL-1ra, IL-6, IL-10, IL-15, IL-18, MCP-1, MIP-1β, and TNFα were determined from the time of vaccination (BL base line) and challenge throughout the critical phase of the disease. Error bars indicate standard deviation. Statistical significance is indicated (* *p*< 0.05, ** *p*<0.01).

### Humoral immune responses after challenge

Protection by the VSV-EBOV has been demonstrated to be primarily mediated by humoral immune responses ^17^ with T cells playing a minimal role ^18^. Therefore, we focused on investigating humoral immune responses. Analysis of EBOV GP-specific IgM revealed that this antigen-specific response was only measurable in the 10^4^ PFU VSV-EBOV_Kik_ group at 0 DPC (Fig. 3A) likely due to the increased replication kinetic displayed in vaccinated NHPs (Fig. S1C).

**Figure 3.**
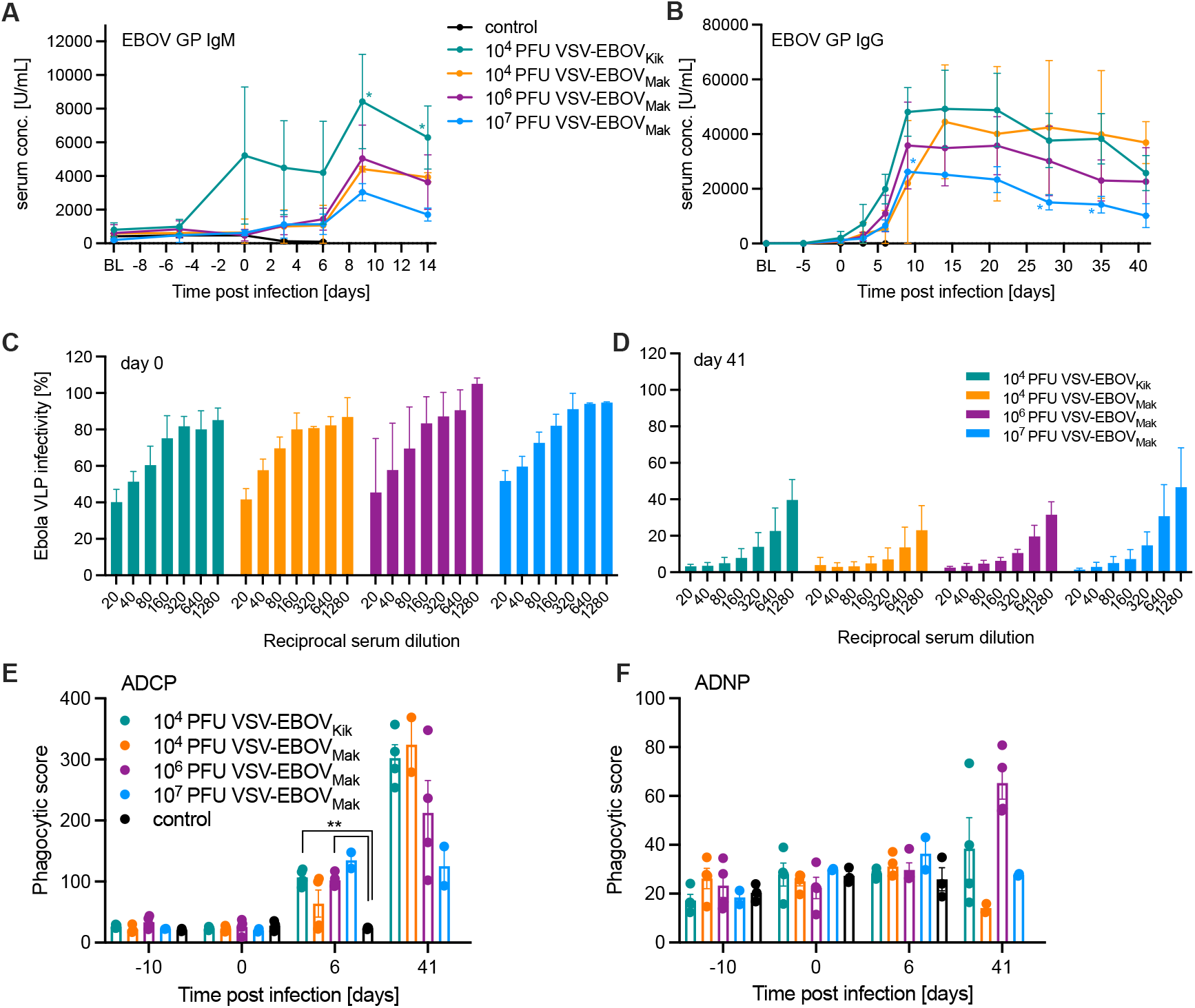
Humoral immune responses in vaccinated and challenged NHPs. EBOV-Makona GP-specific **(A)** IgM and **(B)** IgG levels after vaccination and challenge in serum collected at exam days and the time of euthanasia. Ebola virus-like particle (VLP) neutralizing titers are presented **(C)** at the time of challenge (day 0), and **(D)** at the end of the study (day 41). **(E)** Antibody-dependent cellular phagocytosis (ADCP) and **(F)** antibody-dependent neutrophil phagocytosis (ADNP) at selected time points throughout the study. Error bars indicate standard deviation. Statistical significance is indicated (* *p*< 0.05).

However, EBOV GP-specific IgM levels increased for all vaccinated groups after challenge (Fig. 3A). Similarly, EBOV GP-specific IgG levels were highest in the 10^4^ PFU VSV-EBOV_Kik_ group until 21 DPC, when the two surviving NHPs in the 10^4^ PFU VSV-EBOV_Mak_ group showed higher albeit not statistically significant higher amounts (Fig. 3B). All vaccinated groups showed an increase of EBOV GP-specific IgM and IgG after 0 DPC perhaps indicating a boost effect by the EBOV replication after challenge (Fig. 3A, B). We also determined the neutralizing titers in the serum of the vaccinated NHPs at the time of challenge (0 DPC) and in the surviving animals when the study was terminated (41 DPC) utilizing the EBOV-like particle (VLP) system expressing GFP ^19^. Interestingly, on 0 DPC NHPs in the 10^6^ PFU VSV-EBOV_Mak_ group had the lowest neutralizing titers compared to the other vaccinated groups albeit not significant (Fig. 3C). Reminiscent of the EBOV GP-specific IgG response, the EBOV challenge likely boosted the neutralizing response resulting in 50% reduction of the VLP infection at serum titers >1:640 for all surviving NHPs (Fig. 3D). The neutralizing titers at 41 DPC were not significantly different between the groups, however, it appears that the two surviving NHPs in the 10^4^ PFU VSV-EBOV_Mak_ group have the highest levels (Fig. 3D) matching the highest levels in EBOV GP-specific IgG at that time (Fig. 3B) likely caused by the EBOV challenge serving as a boost of the immune response.

Fc effector function analysis of serum revealed significantly higher antibody-dependent cellular phagocytosis (ADCP) activity for the 10^4^ PFU VSV-EBOV_Kik_ and 10^6^ PFU VSV-EBOV_Mak_ compared to the controls (Fig. 3E). Interestingly, the 2 NHP survivors in the 10^4^ PFU VSV-EBOV_Mak_ group had higher ADCP levels to a similar degree than the other survivors at 6 DPC. Antibody-dependent neutrophil phagocytosis (ADNP) analysis did not result in significant differences at any analyzed time point, however, demonstrated an increase in activity mainly in the survivors vaccinated with 10^6^ PFU VSV-EBOV_Mak_ (Fig. 3F). Interestingly, analysis of antibody-dependent complement deposition (ADCD) revealed a significant difference between the groups achieving uniform protection on 6 DPC (Fig. S3A); like the ADCP results for the antibody-dependent 10^4^ PFU VSV-EBOV_Mak_ group, the 2 NHP survivors had higher levels than the 2 NHPs succumbing to disease in this group (Fig. S3A). Finally, antibody-dependent natural killer cell activation (ADNKA) did not result in any significant fining at any analyzed time point (Fig. S3B).

## Discussion

EVD is listed as one of the WHO’s priority diseases for research and development in emergency events ^20^ despite the availability of licensed vaccines and therapeutics. The approved drugs are monoclonal antibodies and while very potent, they are also vulnerable to be rendered ineffective should the target protein acquire mutations in the antibody binding site. Vaccines are still of great need as the elicited humoral immunity is broader and less vulnerable to a complete loss of protection by viral mutations. The goal is to prevent outbreaks strengthening local, regional and global public health. In this study we emphasized the emergency aspect of the VSV-based EBOV vaccine by investigating the kinetics of protection in a dose de-escalation scenario simulating a potential “dilution of vaccine” to achieve more doses per vial should vaccine shortages occur during an outbreak.

Vaccination with 1×10^6^ PFU of the VSV-EBOV_Mak_ (10-fold reduction from standard dose) resulted in uniform protection with transient mild disease in NHPs infected with a lethal dose of the matching challenge virus, EBOV-Makona. In previous studies characterizing the VSV-EBOV vaccines with 28-35 days between vaccination and challenge in NHPs no challenge virus replication was measured indicating “sterile” immunity ^7,17^. However, when the VSV-EBOV dose was lowered, EBOV viremia was detected highlighting the importance of the vaccine dose for “sterile” immunity ^12^. When NHPs were vaccinated with a single high dose (1×10^7^ PFU VSV-EBOV_Kik_) 7 days prior to challenge, EBOV-Makona was not isolated form the blood demonstrating “sterile” and rapid protection in the context of heterologous challenge ^8^. Here, we emphasize that “sterile” protection is lost to homologous challenge even for the 10^6^ PFU VSV-EBOV_Mak_ dose group in a 10-day vaccination scheme . The data indicate that a lower vaccine dose may not be ideal in an outbreak situation as it could potentially result in virus persistence and delayed transmission. Additionally, a recent study in mice showed that Malaria impacts the magnitude of immunity elicited by VSV-EBOV_Kik_ vaccination^21^.The loss of protective efficacy in mice in this study was overcome by increasing the vaccine dose. Malaria and other conditions including HIV infection impact the human immune system impact immune responses to vaccination and infection ^22,23^. The population in EBOV endemic areas is burdened by HIV and Malaria among others, therefore the reduction of the vaccine dose during outbreaks should only be considered in cases of severe vaccine shortage.

Vaccination with 1×10^4^ PFU of the VSV-EBOV_Mak_ vaccine (1,000-fold dilution from standard dose) achieved only 50% survival demonstrating vaccine breakthrough. Surprisingly, the single dose of 1×10^4^ PFU VSV-EBOV_Kik_ was uniformly protective despite the vaccine antigen being mismatched to the heterologous challenge virus. This is unexpected as it has been shown that VSV-based EBOV vaccines are protective against challenge with any EBOV isolate regardless of the origin of the vaccine antigen ^7,8^. In addition, one would expect same level or increased protective efficacy in a homologous challenge setting compared to heterologous challenge. The VSV-EBOV_Kik_ we used in this study is a GMP produced vaccine stock whereas the VSV-EBOV_Mak_ is a laboratory derived vaccine stock. Basic analysis in the laboratory to this end demonstrated that the non-GMP grade VSV-EBOV_Mak_ preparation contained higher amounts of EBOV GP compared to GMP-grade VSV-EBOV_Kik_ (Fig. S4A, B). However, *in vitro* growth kinetics between the vaccines did not reveal significant differences between the VSV-EBOV vaccines (Fig. S4C, D) whereas *in vivo* VSV RNA levels indicate a slight growth advantage of the GMP produced VSV-EBOV_Kik_ vaccine (Fig. S1C). The higher amounts of GP in the non-GMP grade VSV-EBOVMak preparation may have originated from defective interfering particles or be attributed to increased amounts of shed GP. It could also be that the GMP produced vaccine stock performs better due to its higher purity leading to improved immunogenicity. Further analyses are required to decipher this interesting observation.

While contributing to protective efficacy, the neutralizing antibody response elicited by VSV-EBOV vaccination has not been determined to be a correlate of protection, rather the total EBOV GP-specific antibody response has been shown to corelate with protective efficacy in NHPs and humans ^11,12,17,24^. It is interesting to note that the boost of the total EBOV GP-specific IgG by the EBOV challenge is higher in the lower dose vaccine groups compared to the standard high dose vaccinated NHPs. This correlates with the transient mild signs of disease as indicated by the clinical scores even though infectious EBOV could not be isolated from any of the vaccinated NHPs’ blood samples collected throughout the study. Similarly, the presence of EBOV sGP, the secreted GP from EBOV-infected cells during infection, was only detected in the serum of NHPs that presented with viremia and succumbed to disease. While our Fc effector function analysis overall did not achieve significant results, we identified the trend of higher ADCP activity at 6 DPC likely correlating with disease outcome.

In concordance with previous data ^8,16^ we observed high levels of cytokines and chemokines at the time of euthanasia 5-8 DPC in the five NHPs that succumbed to disease (Fig. 3). The upregulation of these cytokines has also been described for human fatal EVD cases ^25^. Similarly increased levels of IL-1ra, IL-10, and MCP-1 were measured here in the VSV-EBOV_Kik_ group and have been reported one day after high dose vaccination with VSV-EBOV_Kik_ in humans highlighting the vaccine’s ability to quickly induce a strong innate immune response validating our data ^26^.

This study has limitations such as the lack of T cell analysis which was omitted because previous studies demonstrated only a limited role of T cells in VSV-EBOV-mediated protection^17,18^. Another limitation may be related to the differences in product quality of the two VSV vaccines used in this study. Unfortunately, GMP grade material was not available for VSV-EBOV_Mak_. Nevertheless, our study design focused on comparing both vaccines in a rapid protection scenario using the most recent outbreak strain available to us, EBOV-Makona.

Overall, our study highlights that rapid vaccination with reduced doses still leads to protection but at the cost of “sterile” immunity. This raises concerns regarding EBOV persistence and potential downstream transmission. Therefore, lower vaccine doses should only be considered for ring vaccination in cases of severe vaccine shortage.

## Acknowledgements

We are grateful to all members of the Rocky Mountain Veterinary Branch, National Institute of Allergy and Infectious Diseases (NIAID), National Institutes of Health (NIH), particularly the animal caretakers, for their support of this study. This research was supported by the Intramural Research Program of the National Institutes of Health (NIH) (ZIA AI001254 to AM; ZIA AI001089 to HF). The contributions of the NIH authors are considered Works of the United States Government. The findings and conclusions presented in this paper are those of the authors and do not necessarily reflect the views of the NIH or the U.S. Department of Health and Human Services.

## Author contributions

A.M. and H.F. conceived the idea, designed the study, and secured funding. A.M., W.F., A.J.G., F.F., and P.W.H. conducted the animal study. A.M., W.F., A.J.G., K.L.O., E.R.W., and K.S. processed samples, performed assays, and analyzed the data. A.M. and H.F. wrote the manuscript with input from all authors. All authors approved the manuscript.

## Data availability

Data supporting the findings of this study are available at FIGSHARE LINK

## STAR Methods

### Ethics statement

All work involving EBOV was performed in the maximum containment laboratory (MCL) at the Rocky Mountain Laboratories (RML), Division of Intramural Research, National Institute of Allergy and Infectious Diseases, National Institutes of Health (NIH). All procedures followed RML Institutional Biosafety Committee (IBC)-approved standard operating procedures (SOPs). RML is an Association for Assessment and Accreditation of Laboratory Animal Care International (AAALAC)-accredited institution. Animal work was performed in strict accordance with the recommendations described in the Guide for the Care and Use of Laboratory Animals of the NIH, the Office of Animal Welfare and the Animal Welfare Act, United States Department of Agriculture. This study was approved by the RML Animal Care and Use Committee (ACUC), and all procedures were conducted on anesthetized animals by trained personnel under the supervision of board-certified clinical veterinarians. The animals were observed at least twice daily for clinical signs of disease according to a RML ACUC-approved scoring sheet and humanely euthanized when they reached endpoint criteria.

Animals were housed in adjoining individual primate cages that enabled social interactions, under controlled conditions of humidity, temperature, and light (12 hours light - dark cycles). Food and water were available *ad libitum*. Animals were monitored and fed commercial monkey chow, treats, and fruit at least twice a day by trained personnel. Environmental enrichment consisted of commercial toys, music, video and social interaction. All efforts were made to ameliorate animal welfare and minimize animal suffering in accordance with the Weatherall report on the use of nonhuman primates in research (https://royalsociety.org/policy/publications/2006/weatherall-report/).

### Vaccine vectors

Previously described VSV-based vaccine vectors expressing the EBOV-Makona GP, VSV-EBOV_Mak_ ^12^, or EBOV-Kikwit GP, VSV-EBOVKik ^8^, and VSV-MARV ^27,28^ were used in this study. GMP-grade VSV-EBOV_Kik_ was described previously^14^. NHPs were IM-vaccinated with the indicated doses of VSV-EBOV_Mak_, 1×10^4^ PFU VSV-EBOV_Kik_, or 1x 10^7^ PFU VSV-MARV (control); all doses were exactly right as confirmed by backtitration.

### Challenge virus

EBOV-Makona Guinea C07 was used as challenge virus in this study. The virus was propagated on VeroE6 cells (mycoplasma negative), titered via plaque (PFU) and median tissue culture infectious doses (TCID_50_) assay on VeroE6 cells, and stored in liquid nitrogen (EBOV-Makona, passage 2)^16^. Ten days after vaccination, NHPs were IM-infected with 1×10^4^ TCID_50_ of this virus in 1 ml. Inoculum titer was confirmed by backtitration (1.1×10^4^ TCID_50_/ml).

### NHP study design

Eighteen female cynomolgus macaques, 2.5-3.5 years of age and 2.5-3.3 kg at the time of vaccination, were used in this study. The animals were randomly divided into 5 study groups as outlined in Fig. S1A. All animals received a 1ml IM injection for vaccination into 2 sites in the caudal thighs containing either 1×10^7^ PFU VSV-EBOV_Mak_ (n=2), 1×10^6^ PFU VSV-EBOV_Mak_ (n=4), 1×10^4^ PFU VSV-EBOV_Mak_ (n=4), 1×10^4^ PFU VSV-EBOV_Kik_ (n=4), or 1x 10^7^ PFU VSV-MARV (control; n=4). On day -7, one of the control animals died due to anesthetic complications from a previously unidentified cardiac condition. Upon necropsy, the animal was found to have a congenital heart defect with a thickened ventricular wall. This animal was removed from the study. All remaining animals were challenged IM on day 0 with a lethal dose of 1×10^4^ TCID_50_ (1,000 LD_50_) EBOV-Makona into 2 sites in the caudal thighs as described previously ^8,12^. Physical examinations and blood draws were performed as outlined in Fig. S1B on days -10, -9, -7, -5, 0 (challenge day), 3, 6, 9, 14, 21, 28, and 35 and at euthanasia (day 41 for survivors; humane endpoint for non-survivors). Following euthanasia, a necropsy was performed, and samples of key tissues (liver, spleen, and lymph nodes) were collected for virologic and pathological analysis.

### Hematology and serum chemistries

Blood cell counts were determined from EDTA blood with the IDEXX ProCyte DX analyzer (IDEXX Laboratories, Westbrook, ME). Serum biochemistry (including AST, ALP and BUN) was analyzed using the Piccolo Xpress Chemistry Analyzer and Piccolo General Chemistry 13 Panel discs (Abaxis, Union City, CA).

### VSV RNA, EBOV loads and EBOV sGP

VSV RNA copy numbers in EDTA blood samples prior to challenge were determined using a previously described RT-qPCR assay specific to the VSV nucleoprotein ^14^. EBOV loads in macaque EDTA blood and tissue samples were determined on VeroE6 cells (mycoplasma negative) using a median tissue culture infectious doses (TCID_50_) assay as previously described ^16^. Titers were calculated using the Reed-Muench method ^29^. EBOV sGP levels in the serum of challenged NHPs were determined using the previously established ELISA method in the lab ^15^.

### Assessment of humoral immune responses

Post-challenge NHP sera were inactivated by **γ**-irradiation (4 MRad), a well-established method with minimal impact on serum antibody binding ^30,31^, and removed from the MCL according to SOPs approved by the RML IBC. Titers for EBOV GP-specific IgM and IgG were determined using ELISA kits based on the recombinant soluble EBOV-Makona GPΔTM (Alpha Diagnostics, San Antonio, TX). For this, serum samples were diluted 1:400 (IgM), 1:500 (pre-challenge IgG) or 1:2,000 (post-challenge IgG). The ELISA was performed, and titers were calculated as per manufacturer”s instructions. Neutralizing antibody titers were determined by Ebola VLP neutralization as previously described ^19^.

Fc effector function assays using the EBOV-Makona GPΔTM (Alpha Diagnostics) were conducted as previously described ^32^.

### Serum cytokine levels

Post-challenge macaque sera were inactivated by γ-irradiation (4 MRad) and removed from the MCL according to a SOP approved by the RML IBC. Levels of IFNα were determined using the cynomolgus/rhesus IFNα ELISA Kit (PBL Assay Science) following the manufacturer’s instructions. Serum samples were diluted 1:2 in serum matrix for analysis with Milliplex Non-Human Primate Magnetic Bead Panel as per manufacturer”s instructions (Millipore Corporation). Concentrations for analytes IL-1ra, IL-6, IL-10, IL-15, IL-18, MCP-1, MIP-1β, and TNFα were determined using the Bio-Plex 200 system (BioRad Laboratories, Hercules, CA).

### VSV Western blot analysis and growth kinetics

VSV stock samples were mixed 1:1 (*v/v*) with sodium dodecyl sulfate-polyacrylamide (SDS) buffer + 20% β-mercaptoethanol and heated for 10 minutes at 99 °C. Protein separation on SDS-PAGE was carried out using mini-PROTEAN TGX pre-cast gels (Bio-Rad Laboratories). The proteins were then transferred onto a Trans-Blot polyvinylidene difluoride membrane (Bio-Rad Laboratories). The membrane was blocked with 3% nonfat milk powder in PBS and 0.1% Tween (Sigma-Aldrich, St. Louis, MO, USA) for 1 hour at RT. The membranes were incubated with the anti-EBOV GP (ZGP 12/1.1, 1:10,000; a king gift from Dr. Ayato Takada, Hakkaido University, Japan) and the anti-VSV M (clone 23H12, 1:1000; Kerafast, Newark, CA) primary antibodies for 1 hour at RT. The membranes were subsequently stained with an anti-mouse horse-radish peroxidase (HRP)-labeled secondary antibody at 1:20,000 (Jackson ImmunoResearch Laboratories, West Grove, PA) for 1 hour at RT and imaged with SuperSignal West Pico PLUS chemiluminescent substrate (Thermo Fisher Scientific, Waltham, MA, USA) on an Invitrogen iBright FL1500 imaging system (Thermo Fisher Scientific).

### Statistical analysis

Statistical analysis was performed in Prism 9 (GraphPad, Boston, MA). Data presented in Figs. 3, 4, S1C, and S3C were examined using two-way ANOVA with Tukey’s multiple comparison to evaluate statistical significance at all timepoints between all groups. Significant differences in the survival curves shown in Fig. 1A were determined performing Log-Rank analysis. Statistical significance is indicated as ** *p*<0.01, and * *p*<0.05.

## Supplemental information

**Figure S1.**
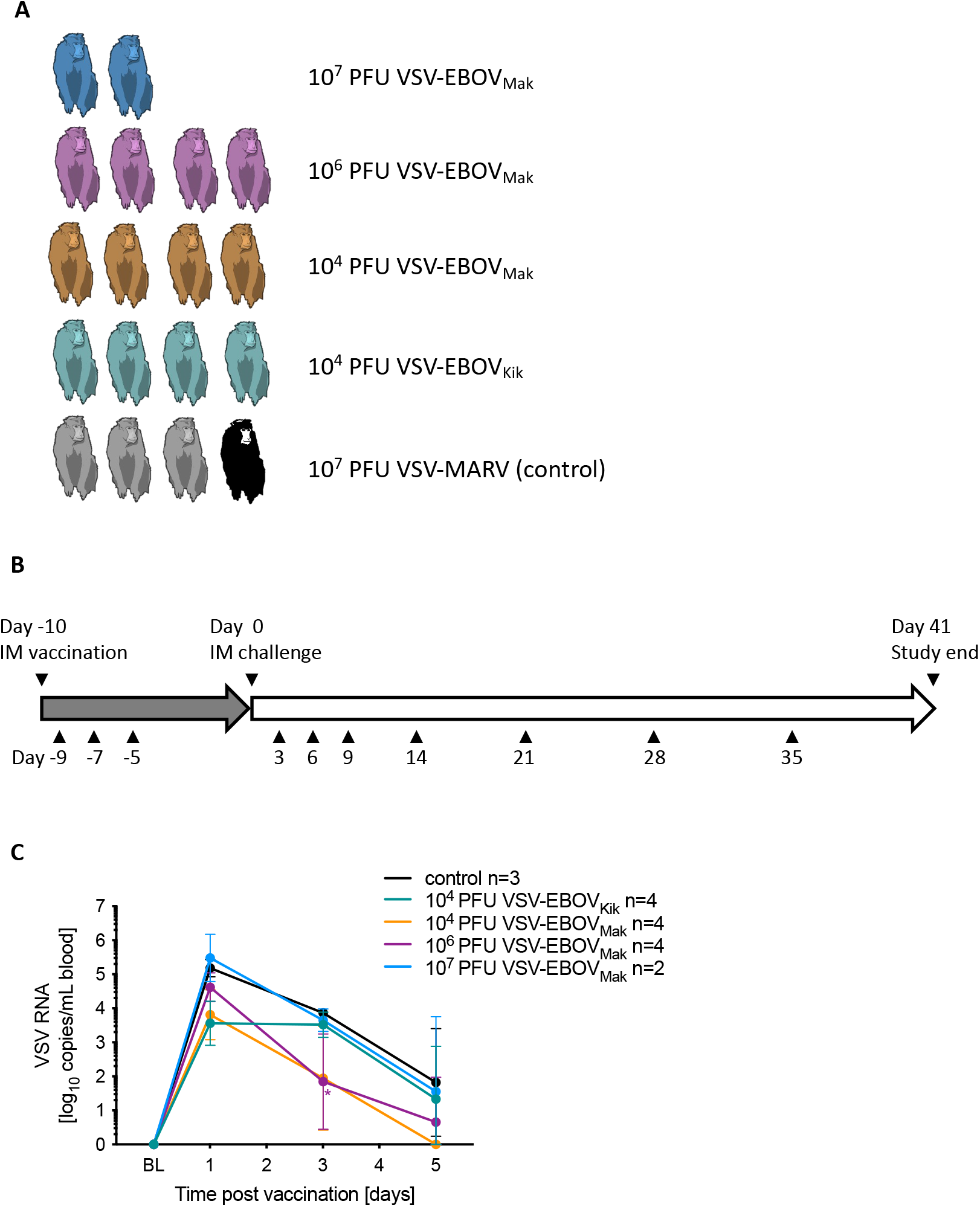
Study design and VSV RNA circulation after vaccination. **(A)** Study groups are depicted. The animal in the control group (gray) that died due to anesthetic complications is indicated in black. **(B)** Vaccination (gray arrow) and challenge schedule (open arrow) for the study. Triangles indicate physical exams including blood draw. **(C)** VSV RNA in the blood of vaccinated NHPs before challenge. BL base line. Statistical significance is indicated (* *p*< 0.05).

**Figure S2.**
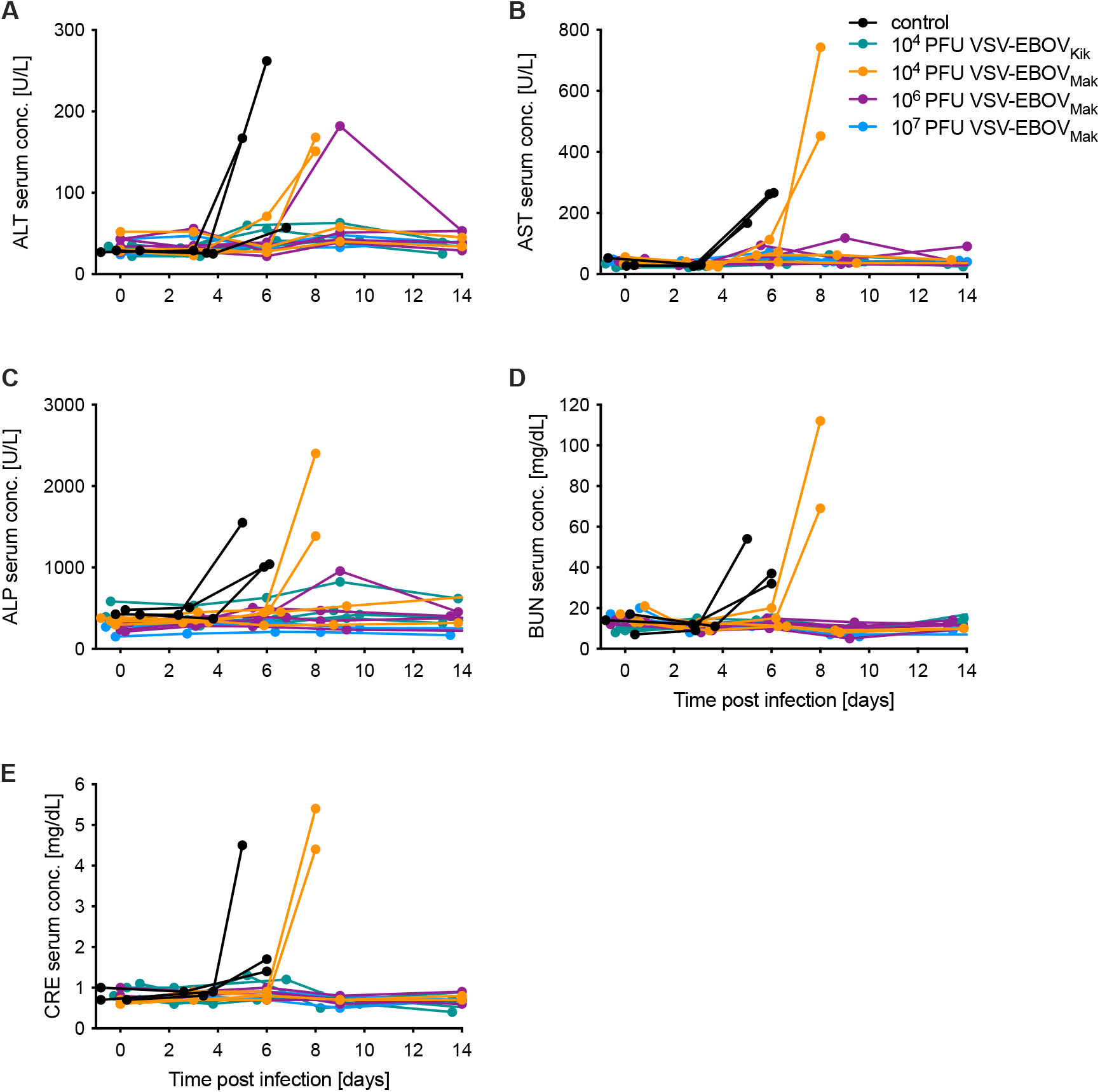
Serum chemistry after EBOV infection in NHPs. The levels of **(A)** alanine aminotransferase (ALP), **(B)** aspartate aminotransferase (AST), **(C)** alkaline phosphatase (ALP), **(D)** blood urea nitrogen (BUN), and **(E)** creatinine (CRE) were determined from serum samples collected after challenge.

**Figure S3.**
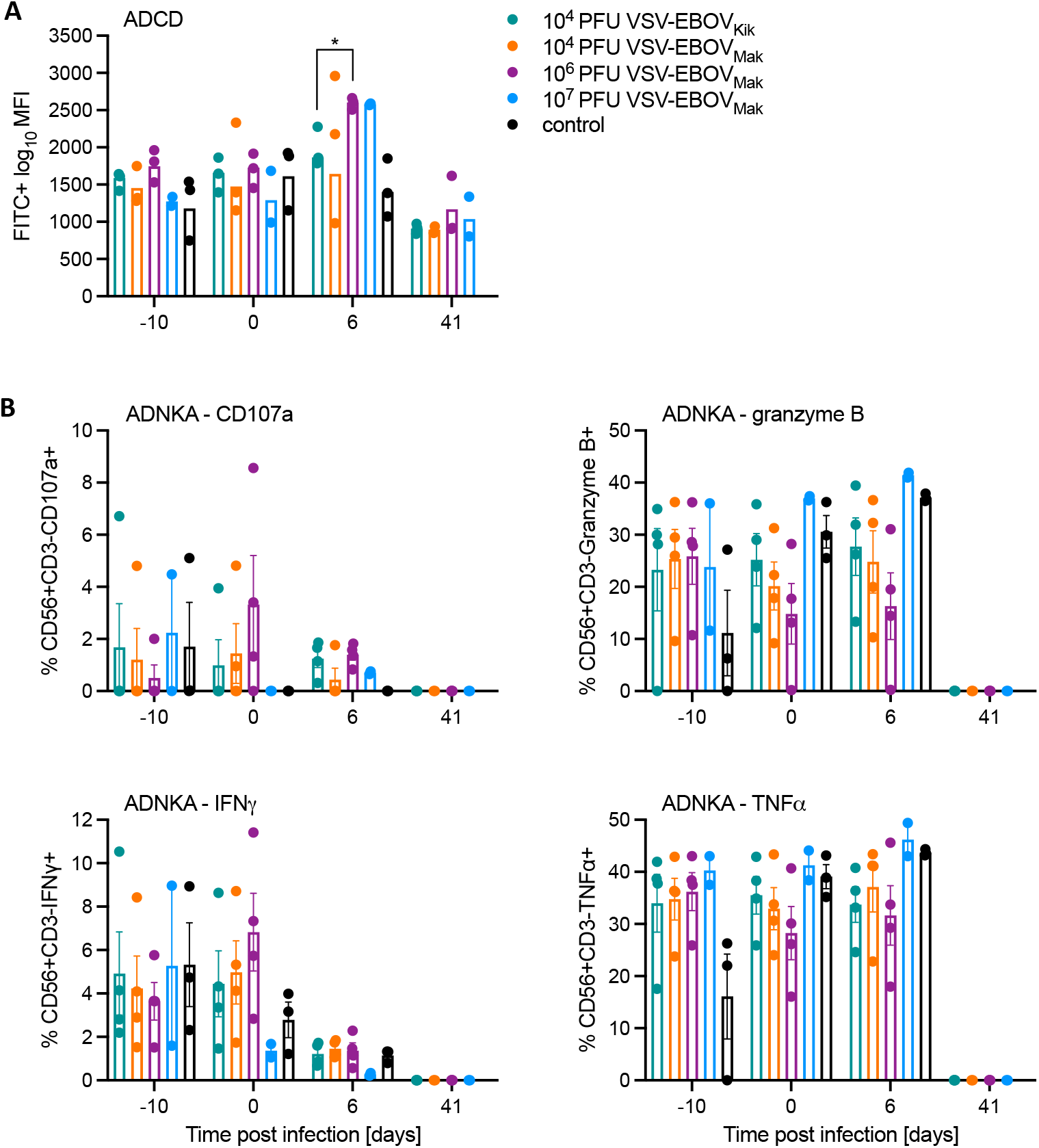
Fc effort function analysis of serum. Levels of **(A)** antibody-dependent complement deposition (ADCD) and **(B)** antibody-dependent natural killer cell activation (ADNKA) were determined at selected time points. Statistical significance is indicated (* *p*< 0.05).

**Figure S4.**
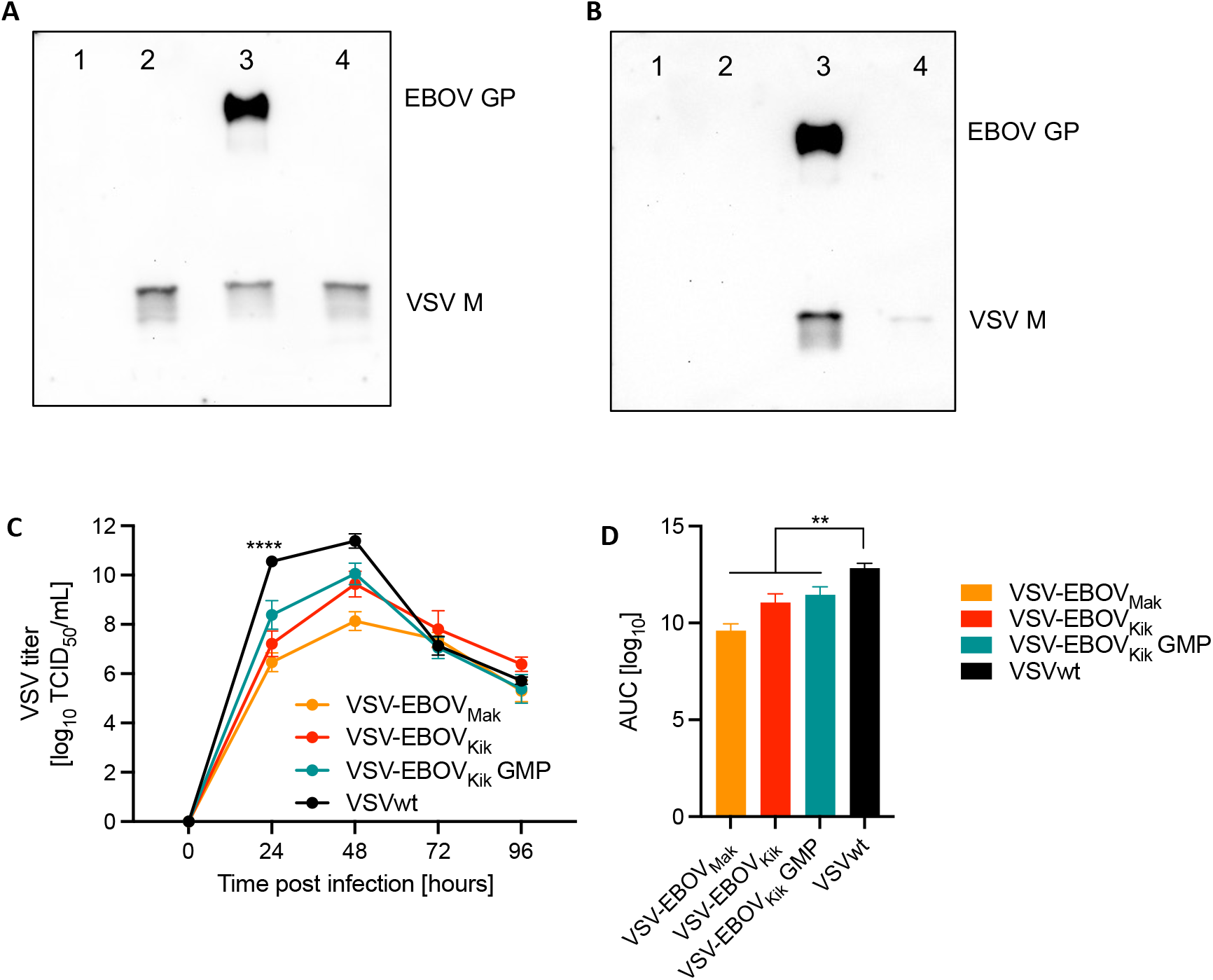
VSV vaccine characterization *in vitro*. Western blot analysis of **(A)** virus stock and **(B)** 1x 10^4^ PFU matched titers of VSVs. Lane 1: uninfected Vero E6 cell supernatant; lane 2: VSV wildtype (non-GMP-grade); lane 3: VSV-EBOV_Mak_ (lane 3; non-GMP-grade); lane 4: VSV-EBOV_Kik_ (lane 4; GMP-grade). Mouse monoclonal antibodies specific to EBOV GP and VSV M were used. **(C)** VSV growth kinetics at a MOI=0.001 on Vero E6 cells. Statistical significance is indicated (* *p*< 0.05).

